# Local postural changes elicit extensive and diverse skin stretch around joints, on the trunk, and the face

**DOI:** 10.1101/2024.10.21.619390

**Authors:** Mia Rupani, Luke D Cleland, Hannes P Saal

**Affiliations:** Active Touch Laboratory, School of Psychology, University of Sheffield, Sheffield, UK; Insigneo Institute for in silico Medicine, University of Sheffield, Sheffield, UK

## Abstract

Skin stretch, induced by bodily movements, offers a potential source of information about the conformation of the body that can be transmitted to the brain via stretch-sensitive mechanoreceptive neurons. While previous studies have primarily focused on skin stretch directly at joints, here we investigate the extent and complexity of natural skin stretch across various body regions, including the face and trunk. We used a quad-camera setup to image large ink-based speckle patterns stamped on participants’ skin and calculated the resulting stretch patterns on a millimeter scale during a range of natural poses. We observed that skin stretch associated with joint movement extends far beyond the joint itself, with knee flexion inducing stretch on the upper thigh. Large and uniform stretch patterns were found across the trunk, covering considerable portions of the skin. The face exhibited highly complex and non-uniform stretch patterns, potentially contributing to our capacity to control fine facial movements in the absence of traditional proprioceptors. Importantly, all regions demonstrated skin stretch in excess of mechanoreceptive thresholds, suggesting that behaviorally relevant skin stretch can occur anywhere on the body. These signals might provide the brain with valuable information about body state and conformation, potentially supplementing or even surpassing the capabilities of traditional proprioception.

## Introduction

The human sense of touch plays a critical role in our interaction with the environment and in learning about our own bodies. Skin stretch, a fundamental component of touch, provides us with rich sensory information about both of these aspects. When we contact an object, even a slight slip (Delhaye et al., 2016) can trigger a surge of neural activity (Delhaye et al., 2021). Perceptually we are also exquisitely sensitive to stretch when applied to the skin passively (Olausson et al., 1998). Beyond external stimuli, skin stretch also contributes to proprioception, our sense of body position and movement. As we flex our joints, such as the fingers or knees, the resulting skin deformation provides valuable cues that complement the signals from traditional proprioceptors (Edin and Johansson, 1995; Collins and Prochazka, 1996; Collins et al., 2005; Kuling et al., 2016). Indeed, some mechanoreceptors are highly sensitive to skin stretch elicited by joint movements, responding to stretch that causes the skin to elongate by as little as 1% (Edin, 1992).

While the role of skin stretch in joint movement has been explored, a critical question remains: how far does this influence extend beyond the immediate vicinity of joints? Moreover, how does skin stretch operate in regions with more complex underlying anatomy, such as the trunk or the face? In the face, specifically, lacking traditional proprioceptors (Cobo et al., 2017), skin stretch has been proposed as a potential source of information for facial proprioception (Staloff and Rafailovitch, 2008; Ito and Ostry, 2010). However, the complex nature of facial movements compared to simpler hinge joints necessitates a deeper understanding of the stretch signal. Existing research on skin stretch has often relied on simplified, one-dimensional measurements of stretch along a specified axis, failing to capture the full complexity of the stretch signal. As the body undergoes three-dimensional conformational changes, the skin, which envelops it, stretches and compresses in two dimensions on its surface. This spatial complexity raises questions about the signals available to the brain and how they might be interpreted to estimate body configuration.

To address these limitations, this study employs a high-resolution measurement approach to map skin stretch across various body regions during static postures. By quantifying the extent and characteristics of skin stretch, we aim to further elucidate its potential role in proprioception and, more generally, the perception of local skin conformation. The findings will lay the groundwork for further research into how the brain integrates and utilizes skin stretch information to arrive at a rich and nuanced understanding of our body in the world.

## Results

We developed a multi-camera setup for gathering high-resolution simultaneous images of random speckle ink patterns applied to the skin across large patches (Fig. 1A, see Methods for details). Correlating the local speckle patterns across multiple cameras allowed us to automatically reconstruct the three-dimensional surface and characterize the shape of complex skin deformation under different poses (Fig. 1B). From these measures we calculated the principal stretch magnitudes and their directions (see Methods) at a resolution of around 1 mm^2^, yielding detailed stretch maps across the full patch (Fig. 1C). We also characterized how the skin deformed in the direction orthogonal to the principal stretch, where it could also stretch or sometimes compress (Fig. 1D). Using this general processing pipeline, we measured stretch induced by different body postures at a variety of skin regions, such as at and around joints, on the trunk, as well as on the face (Fig. 1E) for patches up to 369 cm^2^ in size across 15 participants.

**Figure 1.**
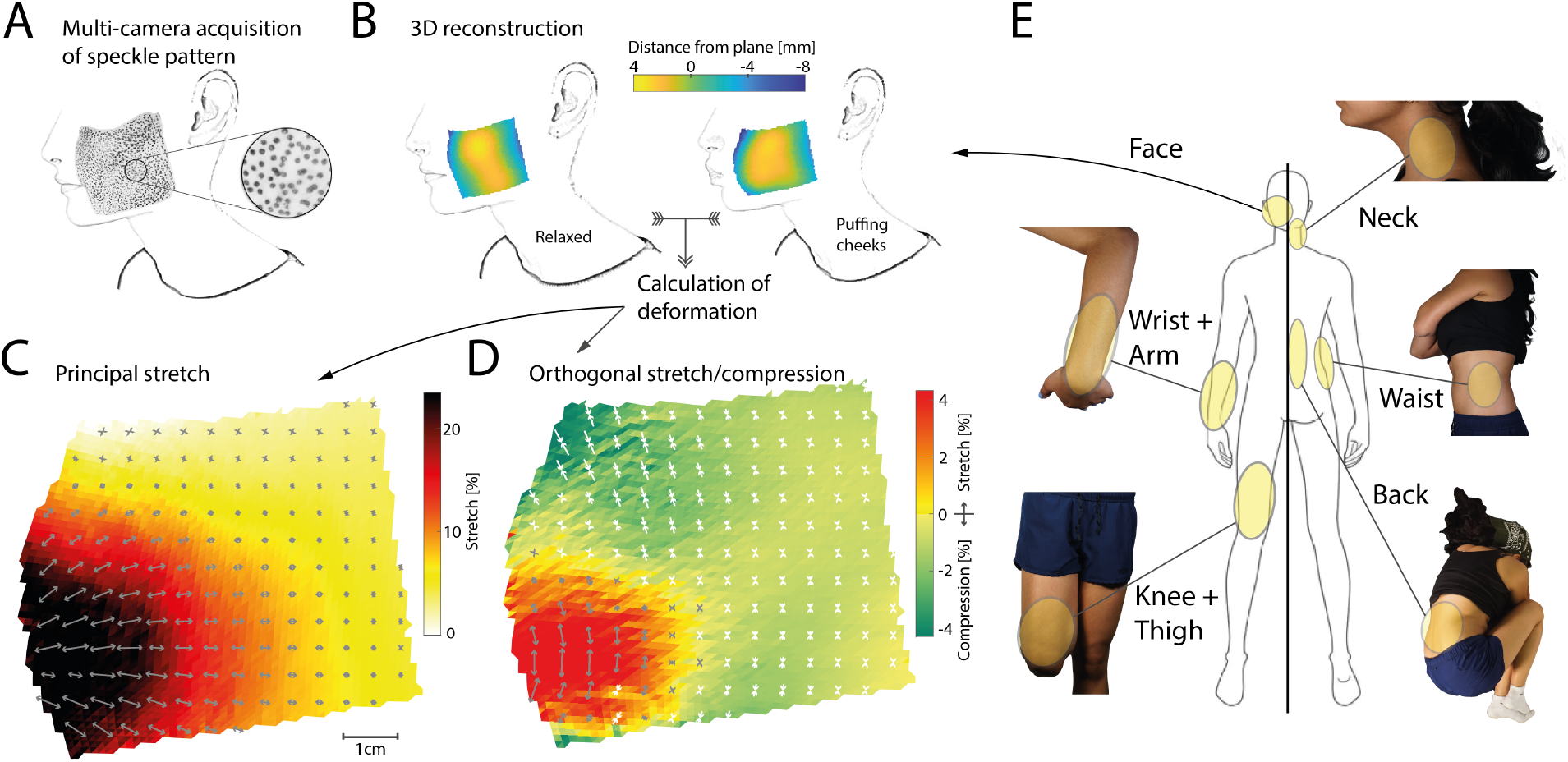
Data acquisition framework and targeted body regions. **A**. An ink-based random dot speckle pattern was applied to the targeted skin region and imaged through up to four high-resolution cameras simultaneously. The example shown here is a roughly 8×8 cm patch applied to the cheek of a participant. Individual speckle dots are clearly visible in the inset. The original photograph has been digitally converted to outlines with some features removed. **B**. Local speckle patterns are correlated across two or more cameras using digital image correlation to obtain a three-dimensional reconstruction of the imaged skin surface patch. In the examples shown, a two-dimensional plane has been fit to the reconstructed three-dimensional surface, with colours denoting the distance of each local patch from that plane, illustrating the shape of the cheek area. Left: A relaxed pose, serving as the baseline for stretch calculations. Right: The participant is puffing their cheeks, which results in a bulging of the cheek area close to the mouth in the 3D reconstruction. Considering both reconstructions together allows calculation of skin deformation from the baseline relaxed pose. **C**. Magnitude (coloured shading) and direction (arrows) of the principal stretch, i.e. the stretch along the direction in which it largest, when the participant is puffing their cheeks. Darker shading indicates larger stretch which reaches upwards of 25 % close to the mouth. Arrows indicate the direction of stretch, which is not uniform across the imaged area. All measures are shown projected onto the best-fitting plane, with stretch magnitudes subsampled to 1 mm and directions subsampled to 5 mm resolution for better visibility. **D**. Magnitude and direction of stretch in the direction orthogonal to the principal direction (as shown in C). Yellow/red shading indicates positive stretch, implying that in these regions the skin expands in all directions. Green shading and white arrows indicate compression. Note the generally smaller stretch magnitudes along this secondary axis. **E**. Body regions where stretch values were measured. These include the face, the wrist/arm and knee/thigh regions, as well as the neck, waist, and back.

### Large spatial extent of skin stretch around joints

First, we investigated skin stretch around joints where it has been implicated in proprioception. Prior work had reported that putative stretch-sensitive mechanoreceptors on the upper thigh responded to even slight knee flexion (Edin, 2001), indicating that skin stretch induced by joint movements might extend beyond the immediate joint vicinity. Our imaging analysis of skin stretch from the knee to the upper thigh during maximal knee flexion confirmed this hypothesis. As expected, under maximal knee flexion we observed large stretch in excess of 60% directly at the knee joint, but considerable stretch of 10- 20% at the upper thigh (Fig. 2A). The direction of principal stretch was away from the joint, but angling outwards and inwards to the lateral and medial sides of the thigh respectively (Fig. 2B). As a consequence the central portion of the thigh displayed stretch in both its principal direction and orthogonally, as the skin was pulled bi-directionally. This pattern persisted across multiple participants: stretch was highest directly at the knee, but persisted above 10% even 15-20 cm away (Fig. 2C), likely continuing beyond the measured range. However, skin stretch was not solely determined by distance from the knee, but also varied across thigh locations, with the lateral side of the knee exhibiting smaller stretch magnitudes than the medial side (Fig. 2D).

**Figure 2.**
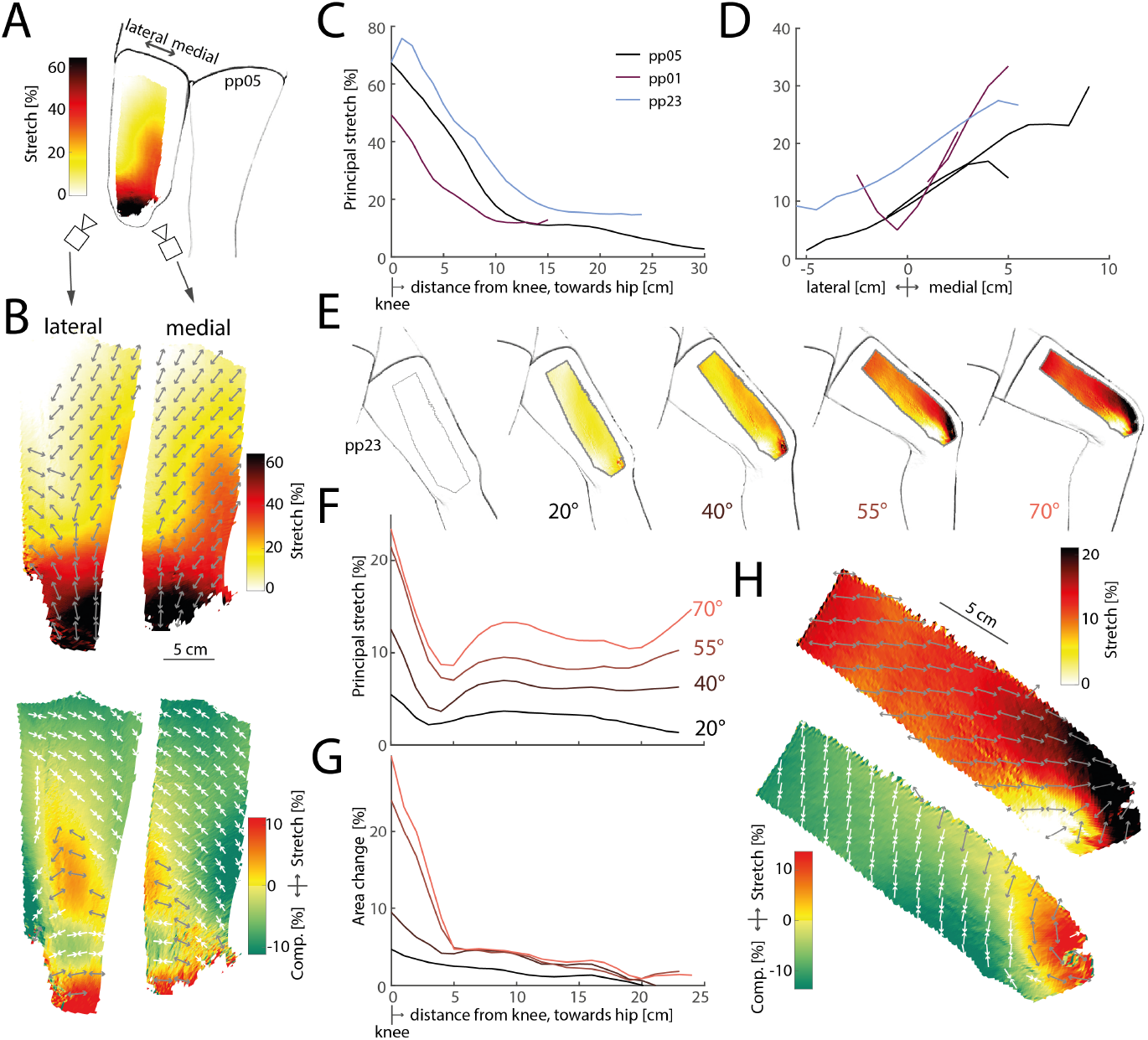
Stretch patterns on knee and thigh skin in response to flexion of the knee. **A**. Principal stretch on the thigh under maximal flexion of the knee. Stretch reaches more than 60% at the knee, but is still around 20% on the upper thigh. **B**. Detailed maps of principal (top row) and orthogonal (bottom row) stretch on the lateral (left column) and medial (right column) thigh. **C**. Average principal stretch from distal (knee) to proximal (upper thigh) for three participants. Stretch is highest at the knee, extends far along the thigh. **D**. Average principal stretch along the lateral/medial axis for the same participants as in C. Stretch is consistently larger on the medial compared to the lateral thigh. **E**. Stretch on the medial thigh when flexing the knee at different angles. Stretch extends far even at small angles. **F**. Principal stretch from distal to proximal along the medial thigh for the same participant as in E at the same knee angles. Stretch at the upper thigh grows systematically with knee angle. **G**. Local skin area change for the same data as in F. Away from the knee, skin area does not grow with knee angle, suggesting that the increased principal stretch observed in F is compensated for by increased compression in the orthogonal direction. **H**. Detailed maps of principal and orthogonal stretch for knee flexion at 70°angle.

We also tested how skin stretch patterns changed with increasing knee joint angles. For one participant, we imaged the medial thigh during joint angles up to 70°. As expected, principal stretch values increased with joint angle. Notably, the large spatial extent of skin stretch all over the thigh was evident even at the smallest angle tested (20°, Fig. 2E). While principal stretch increased monotonically with joint angle at all measured distances, stretch values close to the knee were close to saturation at higher angles, while continuing to increase further away (Fig. 2F). Thus, skin regions away from joints might provide valuable proprioceptive information at higher joint angles that is not available from skin directly at the joint, which might already be maximally extended.

Analysis of the full two-dimensional stretch patterns revealed distinct differences between skin regions near and far from the knee joint. Skin immediately surrounding the joint tended to stretch along both its principal and orthogonal directions, while skin farther away primarily stretched along its principal axis but contracted in the orthogonal direction (see Fig. 2H for an example). Consequently, while total skin area expanded considerably around the knee, regions distant from the joint exhibited moderate or no change in skin area, even at the largest joint angles tested (Fig. 2G). These findings impose constraints on whether stretch-sensitive mechanoreceptors relay the relevant signals: receptors near the joint can be sensitive to stretch in any direction and reliably convey joint angle information, while receptors elsewhere on the thigh would need to be responsive to stretch along the principal axis specifically, as the skin is only stretched along this axis.

The general trends observed for the knee were mirrored when measuring stretch on the forearm in response to flexion of the wrist (Fig. 3A,B). Stretch was highest directly at the wrist, but extended onto the forearm and was still measurable even close to the elbow (Fig. 3C). Again, the skin was stretched in all direction close to the joint, but only stretched along its principal direction further distally on the forearm, while slightly compressing in the orthogonal direction (Fig. 3D). As on the knee, the principal direction was not perfectly aligned with the proximal-distal axis along the limb, but instead angled sideways (e.g. medially in the example shown in (Fig. 3B).

**Figure 3.**
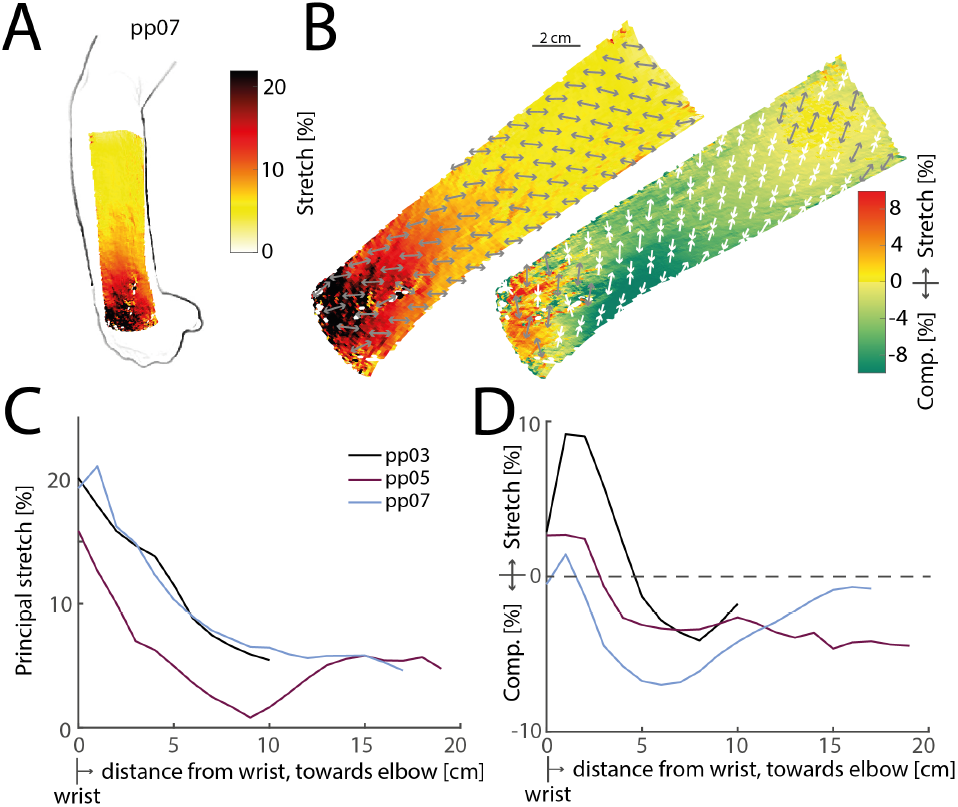
Stretch on the forearm due to wrist flexion. **A**. Illustration of principal stretch measured on a participant’s forearm as the wrist is maximally flexed. **B**. Detailed maps of principal (left) and orthogonal (bottom row) stretch for the same participant shown in A. **C**. Average principal stretch at distally increasing distances from the wrist towards the elbow for three participants. Stretch is highest at the wrist, but extends all the way towards the elbow. **D**. Same as in C, but for stretch orthogonal to the principal direction. The skin is stretched close to the wrist, but compresses in this direction in the middle of the forearm.

In summary, skin stretch due to joint movements is not limited to skin directly at the joint itself, but extends across the involved limbs.

### Uniform and large skin stretch on the trunk

Next, we investigated skin stretch in body regions surrounding the trunk, including the neck, back, and waist. Unlike hinge joints such as the knee, the neck contains pivot joints, enabling more complex movements. The waist and back are influenced by movements of the spine. Collectively, the trunk region constitutes a large portion of our total body surface, potentially providing strong stretch feedback despite its relatively low innervation density (Corniani and Saal, 2020).

We found that tilting the head sideways to induce stretch on the neck (Fig. 4A), curling up in a fetal position to stretch skin on the back (Fig. 4B), and bending sideways while standing to stretch the side of the waist (Fig. 4C) consistently induced large, relatively uniform skin stretch that extended beyond the boundaries of tracked regions for all participants. Even the lowest stretch values measured approached 10%, increasing in some cases beyond 50%, and averaging 22% on the neck, 23% on the back, and 29% on the waist across all participants (Fig. 4D). Orthogonal to the principal stretch, we measured a slight compression of the skin, averaging between -2% and -3% on all three sites. A notable exception was the skin directly above the spine, which expanded in all directions and exhibited stretch both longitudinally along the spine as well as laterally, with individual vertebrae visible (see Fig. 4E, right panel).

**Figure 4.**
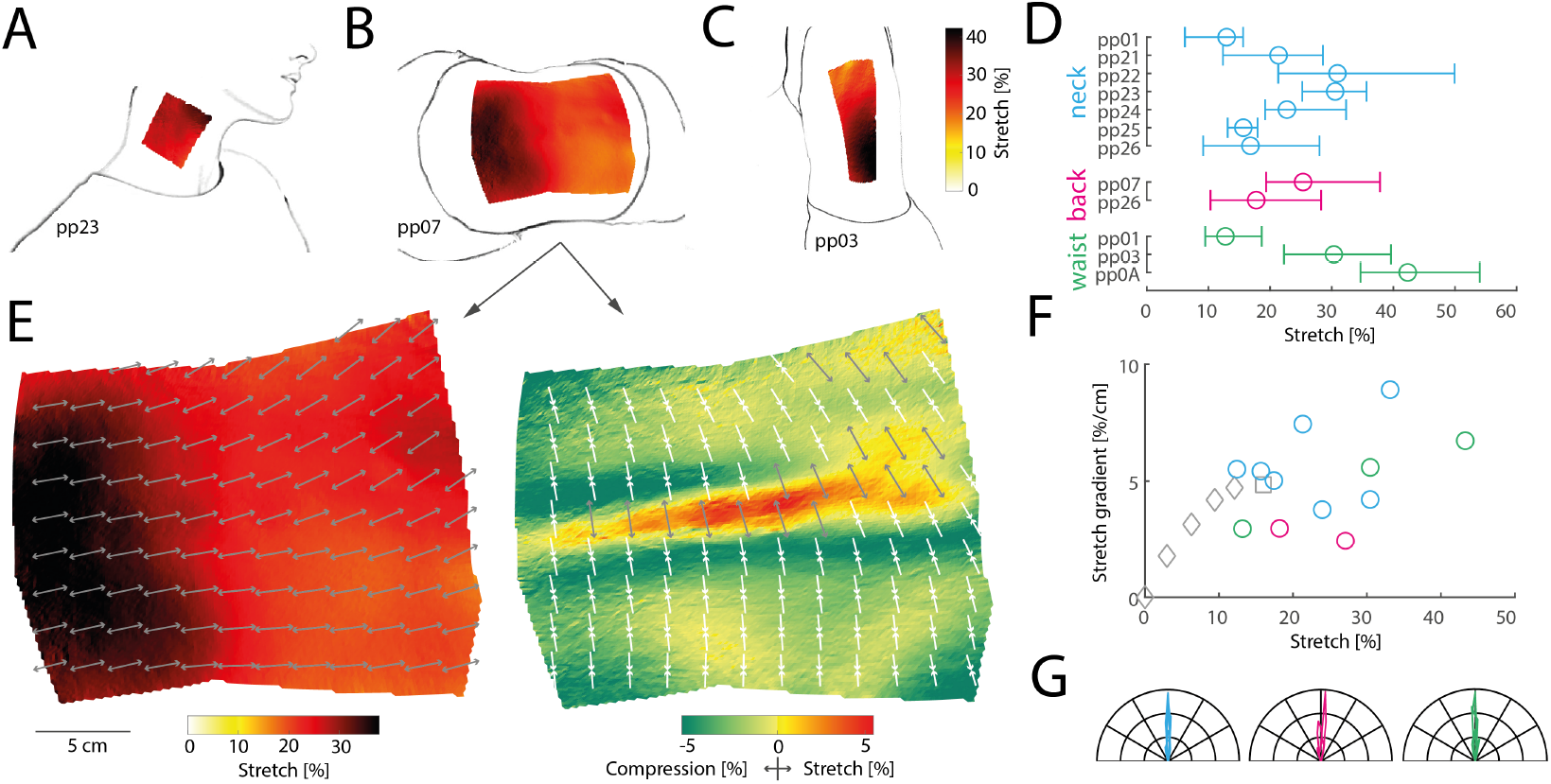
Stretch on the body’s trunk. **A**,**B**,**C**. Illustrations of principal stretch measured on a participant’s neck as the head is bent sideways (A), on the back as the body curls up in the fetal position (B), and on the waist as the body bends sideways (C). **D**. Average stretch, and 5th and 95th percentiles for individual participants measured at the neck (blue), back (pink), and waist (green). **E**. Detailed principal stretch (left panel) and orthogonal stretch (right panel) on the back for the same participant as in B. The tracked surface is projected onto the best fitting plane. Arrows indicate the direction of stretch (gray: elongation; white: compression), subsampled and displayed every 2 cm for visibility. The tracked mesh includes 39,139 triangles in total, with an average area of just below 1 mm^2^ and a total tracked area of 369 cm^2^ in the unstretched state. **F**. Average principal stretch versus stretch gradient for all areas on the trunk (colored circles) with data from the knee (in grey; diamonds: increasing knee flexion; square: full knee flexion). **G**. Polar histograms showing the relative frequency of the principal orientation of stretch across all tracked triangles projected onto the best-fitting 2D plane for the neck, back, and waist. Each coloured line denotes a different participant. For each distribution, the average principal stretch orientation has been subtracted from each measured orientation, such that 0°denotes this average, while negative and positive angles show deviations from the average. Stretch is almost entirely uniform for all participants and all skin sites.

Stretch on the trunk was spatially uniform. While some subregions exhibited greater stretch than others, the stretch gradient (change in stretch over distance) averaged around 5%/cm, varying across participants and skin sites but never exceeding 10%/cm (see Fig. 4F, coloured circles). This suggests that stretch on the trunk decays slowly with distance, affecting a large skin area, similar to our observations on the knee (see Fig. 4F, grey symbols). Stretch was also uniform in its principal orientation: the average angular standard deviation was 5.7°on the neck, 9.6°on the back, and 5.2°on the waist, suggesting that the direction of stretch varied little even across large patches of skin on the trunk (Fig. 4G). In summary, skin stretch on the trunk is of large magnitude and spatially extensive, but relatively uniform.

### Spatially complex skin stretch on the face

To explore skin stretch in a region known for its complex movements and lack of traditional proprioceptors, we focused on the cheeks. Facial muscles in this area interact to support facial expressions, speech, and mastication, likely resulting in intricate skin deformations. Previous research has suggested that skin stretch serves as a major feedback signal during facial motor control and learning. We measured skin stretch across four distinct facial poses, highlighting the breadth of facial movements: puffing the cheeks, pouting, opening the mouth widely, and smiling (see Fig. 5A for illustrations and Methods for corresponding action units in the Facial Action Coding System).

**Figure 5.**
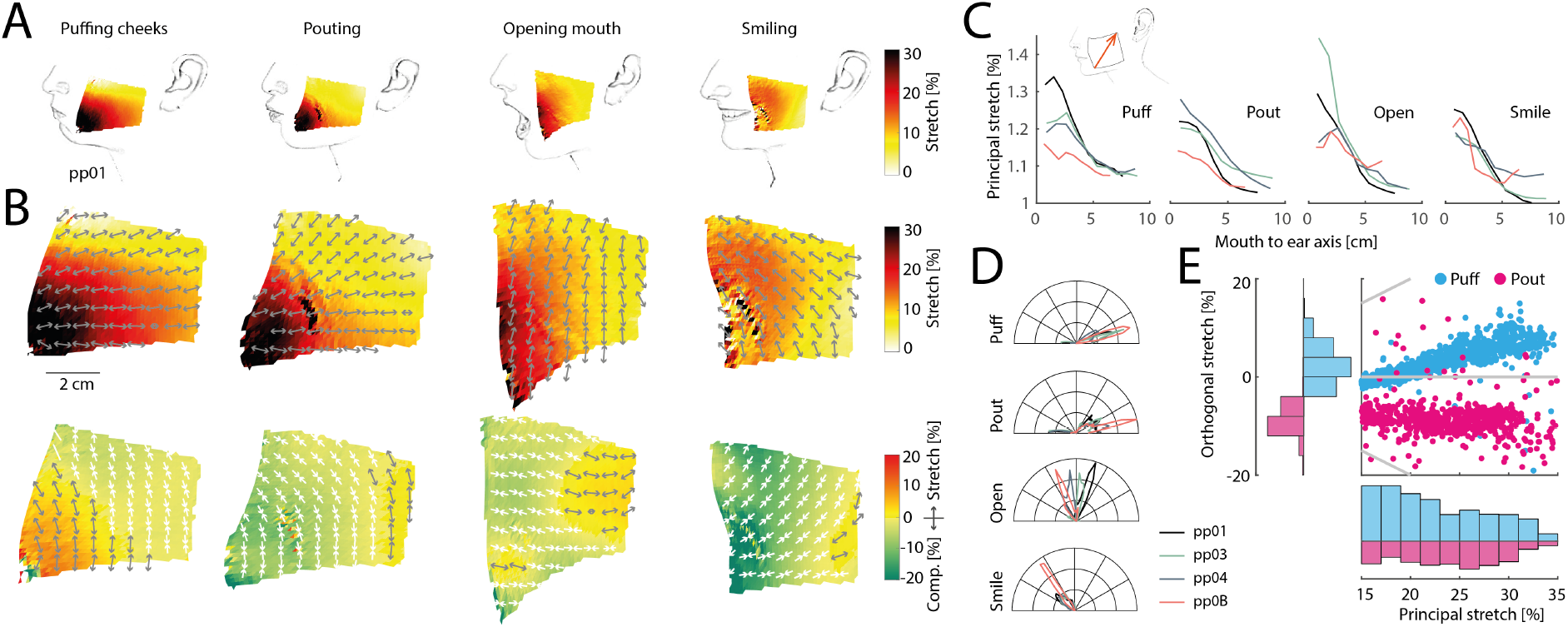
Skin stretch on the face during a variety of facial movements. **A**. Illustration of poses, tracked area on the cheek, and principal stretch for one participant. **B**. Detailed maps of principal (top row) and orthogonal stretch (bottom row) for the same participant as in A. **C**. Stretch for all participants projected onto an axis extending from the lower left corner of the tracked area to the upper right (see illustration in inset), therefore roughly extending from the lower mouth in the direction of the ear. Stretch is highest close to the mouth in all four poses and for all participants and decays towards the ear. **D**. Polar histograms showing the relative frequency of the principal orientation of stretch across all tracked triangles projected onto the best-fitting 2D plane in the unstretched state. Different panels show different poses. Raw distributions are shown for each pose, so that different angular orientations become evident. **E**. Scatter plot (top right) and histograms (left and bottom) of principal and orthogonal stretch for same participants as in A during puffing (blue) and pouting (pink). The extent and distribution of the principal stretch is similar in both conditions, while the orthogonal stretch differs, yielding extension in the puffing condition and compression in the pouting condition.

These poses yielded unique and complex skin stretch patterns compared to other body regions (see Fig. 5B for examples of detailed principal and orthogonal stretch maps). Stretch was generally larger closer to the mouth than laterally on the cheek. Indeed, when projecting stretch values onto an axis leading from the lower corner of the tracked region (located close to the mouth) to the upper lateral corner (towards the ear), stretch was consistently higher closer to the mouth for all participants and poses (Fig. 5C). The orientation of stretch also varied more on the cheek area compared to the larger trunk regions, with average angular standard deviations more than two-fold those recorded on the trunk (13.0° for puffing, 20.8° for pouting, 17.6° for opening the mouth, and 16.1°for smiling, see Fig. 5D). Furthermore, different poses produced distinct distributions of skin stretch orientations, with opening the mouth eliciting principal orientations nearly perpendicular to those during puffing or pouting (see also Fig. 5B, gray arrows in top row, for individual examples). This suggests that distinguishing between these poses based on stretch feedback requires considering both stretch magnitude and orientation.

Finally, we noticed that puffing and pouting yielded similar principal stretch patterns: in both instances, stretch was similar in magnitude, spatial arrangement, and orientation (see Fig. 5B, first two panels on top row). However, these two poses resulted in different three-dimensional cheek geometries. While puffing involves outward bulging of the cheek, pouting leads to a moderate inward contraction. To successfully disambiguate between these two poses the brain might use information about skin stretch in the direction orthogonal to its principal stretch (see Fig. 5B, first two panels on bottom row): while orthogonal skin stretch was positive in the puffing condition, signaling an overall expansion of the skin area driving by the outwards bulging of the cheek skin, orthogonal stretch was negative in the pouting condition, signaling a contraction of the skin along this axis (see scatter plot and histograms in Fig. 5E). This suggests that fully exploiting relevant information in active skin stretch patterns requires mechanoreceptors sensitive to the entire complexity of two-dimensional skin stretch, rather than just coarse measures along a single axis.

In summary, stretch on the face appears more complex compared to other body regions, with most stretch concentrated towards the mouth area.

## Discussion

We measured natural skin stretch induced by different body postures and muscle movements around joints, on multiple trunk regions, and on the face. We found widespread and often complex stretch patterns in all regions tested. Around joints, stretch could be measured far from the joint itself, with knee flexion inducing notable skin stretch as far as the upper thigh, tens of centimeters away. On the trunk, bending of the body led to large regions of uniform stretch that often exceeded 20% and more. Finally, different facial poses elicited complex and varying stretch patterns on the cheek, with the highest stretch values measured close to the mouth.

### Active stretch induced by body movements

Contact with surfaces often induces shear forces and therefore skin stretch, which can be a salient signal about contact properties. For example, the onset of slip during object grasping induces shear waves on the skin (Delhaye et al., 2016, 2021), which can serve as a rapid signal for making corrective actions. During full slip, skin stretch contributes to signaling the direction of the moving object (Seizova-Cajic et al., 2014). Shear forces can also signal the body’s movements relative to its surroundings, such as on the foot sole during walking (Crossland et al., 2022) or on the buttocks when seated during acceleration of a vehicle (Horie et al., 2018).

In contrast to these externally generated skin stretches, here we focused on active skin stretch that is generated by the body itself and that carries information about body conformation. Previous work has mostly focused on skin stretch directly at joints where stretch can be expected to signal joint angle. However, it is likely that relevant stretch signals occur not only in skin directly surrounding the joint, but extend further across the body. For example, flexion of the elbow has previously been shown to stretch skin on the forearm (Maiti et al., 2016) and different ankle postures elicit skin stretch across the foot sole and dorsum, respectively (Smith et al., 2019). In the present work, we therefore took a more expansive approach and considered stretch at a multitude of skin regions. Some of the chosen region have received previous attention: stretch on the back has been investigated using a marker-based setup (Beaudette et al., 2017) as well as optically (Kao et al., 2022, 2023), and small patches of skin on the face have previously been imaged for skin stretch (Staloff et al., 2008). By employing a fully optical method with high spatial resolution, we were able to image skin patches much larger than in previous efforts. As our setup was optimized for imaging large patches of skin with high resolution, this limited the rate at which images could be acquired. We therefore focused on static postures, ignoring dynamic aspects of the stretch signal. However, choosing a different trade-off between resolution and frame rate, or using more advanced cameras would allow recording dynamic information as well (see Kao et al., 2022, 2023, for examples).

### Extensive skin stretch across the body

We found that skin stretch often occurs over extensive patches of skin, therefore potentially providing strong and rich feedback. What are the main causes driving widespread stretch? Mechanically, some joints possess skin folds, such as those on the back of the fingers, that support a large movement range as they unfold and absorb the mechanical deformation locally at the joint. A similar principle is at play in elephant trunks, which also display prominent skin folds (Schulz et al., 2022). However, many human joints do not possess distinctive skin folds and are therefore limited to the skin’s general capacity for stretch, which has been found to be anisotropic due to the skin’s natural tension (Ní Annaidh et al., 2012). Even in the absence of folds, the skin exhibits furrows and ridges, which mechanically support stretching to some extent (Leyva-Mendivil et al., 2015; Corniani et al., 2023), but might cause more widespread stretch further from the joint to fully support large joint movements. The large extent of stretch along the thigh we observed might partially be caused by this effect. Other body movements, such as bending of the trunk, are not supported by hinge joints, but instead deform the body over a larger region. resulting in extensive skin stretch across large parts of the body. Finally, the underlying geometry of muscles and other body parts themselves can change and deform the skin directly above them, without any direct change in the body’s pose. Indeed, tensing of the muscles without any movement of the knee will change the skin conformation of the thigh area especially on the medial side, likely leading to measurable skin stretch patterns without any overt movement of the leg. Thus, skin stretch need not be induced by joint movements per se, but could also respond directly to shape changes by the muscles during activation. Overall, the consequence of these disparate mechanisms appears to be widespread stretch patterns.

### Stretch as a rich multi-dimensional feedback signal

While skin stretch often exhibited a relatively uniform spatial distribution, particularly on the trunk, its complexity varied across body regions. On the face, for instance, stretch patterns were more intricate, varying spatially in both their principal and orthogonal directions. These findings suggest that skin stretch is not a simple one-dimensional signal along a constant axis defined solely by its magnitude, but rather a complex multidimensional phenomenon: the orientation and magnitude of stretch along both its principal and orthogonal directions all provide valuable and independent cues about changes in body shape. For example, the act of puffing the cheeks results in an expansion of the total skin area of the cheek, as measured by stretch along both the principal and orthogonal axes. An expanding cheek area coupled with the surrounding skin not deforming, is only possible if the cheek area deforms in the third dimension, either inward or outwards. Thus, stretch signals can, in some circumstances, provide hints about the three-dimensional shape of body parts. It is likely that such signals are available to and used by the brain, though perhaps in more basic form, when originating from facial skin: skin deformation and its associated cutaneous cues have been implicated in motor learning (Ito and Ostry, 2010) and speech perception (Ito et al., 2009; Franken et al., 2022), We also possess at least partial awareness of and control over our facial expressions (Ciston et al., 2022), which signal not only the emotional category but also their intensity (Chen et al., 2024), serve as salient cues to others during social interactions (Chen et al., 2020), and modulate incoming information from other sensory systems (Susskind et al., 2008).

More generally, stretch feedback might support mechanisms that support body perception beyond proprioception, by supplying information about the state of the skin, sometimes called dermatokinesthesis (Halpern, 1946), about the size and shape of the body, termed somatoperception (Longo et al., 2010; Longo and Haggard, 2012), and perhaps even support representations more classically associated with interoception (Crucianelli and Ehrsson, 2023).

### Neural mechanisms and perceptual consequences

To effectively utilize stretch signals, especially their complex aspects, the nervous system must first transduce them via mechanoreceptive neurons. Recordings from human primary tactile demonstrate that a significant proportion of primary tactile neurons innervating the dorsal skin of the hand (Edin and Abbs, 1991) and thigh (Edin, 2001) are highly sensitive to skin stretch induced by joint movements, even without additional object contact. Notable, stretch responses were observed in neurons with small receptive fields located distant from the joints, such as responses on the upper thigh during knee flexion (Edin, 2001), suggesting widespread skin sensitivity to local stretch patterns. Indeed, careful experiments involving passive skin stretching revealed that reliable responses in some receptors could be elicited by stretch as low as 1% (Edin, 1992, 2004), at least an order of magnitude lower than those reported in the present study. This indicates that stretch on all investigated body parts readily exceeds mechanoreceptive thresholds.

Slowly-adapting type II (SA2) neurons, thought to innervate Ruffini corpuscles (Halata, 1988), are well-known for their sensitivity to skin stretch, responding to both dynamic and static stretch induced by joint movements (Edin and Abbs, 1991). Some SA2 neurons exhibit tonic activity (Chambers et al., 1972), potentially signalling ongoing skin tension, while also modulating their response rate to subtle skin movements in the absence of an external force (Saal et al., 2023). Moreover, stimulation of individual SA2 neurons can often by perceptually detected (Watkins et al., 2022). These characteristics make SA2 neurons particularly well-suited for signalling precise and ongoing information about the skin’s state. However, it is important to note that several other tactile neurons types, including fast adapting ones, also respond to stretch and stretch changes, suggesting a potentially even richer signal.

While not extensively studied, there is evidence that stretch responses are sensitive to the direction of stretch, thereby signaling more intricate aspects of stretch patterns. SA2 neurons have been shown to be sensitive to the direction of stretch, both when the skin is passively stretched (Grigg, 1996) and when it is is recovering mechanically after a stimulus has been applied (Saal et al., 2023). Perceptual experiments have also confirmed our ability to discern the direction of passively applied skin stretch on various body parts (Olausson et al., 1998; Gleeson et al., 2009; Chen et al., 2016). To our knowledge it is currently unknown whether more complex stretch aspects, such as the stretching or compression of the skin orthogonal to the principal stretch direction, are also reflected in neural responses or perceptually available.

Beyond their role in proprioception, skin stretch signals have various perceptual consequences, including changes in thresholds (Beaudette et al., 2017; Smith et al., 2021), two-point discrimination acuity (Cody et al., 2010), tactile distance estimation (Mainka et al., 2023), and localization (Kang and Longo, 2023). These findings, along with our own, suggest a more profound interconnection between touch and proprioception than previously assumed. Similar to how different touch receptors collaborate to signal relevant tactile features (Saal and Bensmaia, 2014; Corniani et al., 2022), information about body state and conformation appears to be distributed across a variety of receptors, encompassing both classical proprioceptive and more traditionally tactile neurons.

## Methods

### Participants

15 participants (7 male, 7 female, 1 non-binary) with a mean age of 24.5 (range: 18-41) years and no known allergy against ink were recruited to take part in the study. Not all participants had all skin regions imaged due to time constraints and limitations of the camera setup; additionally not all trials yielded usable data (see Table 1 for the full list of trials and participants included in this report). All participants provided informed consent prior to the start of data collection. The study protocol was approved by the ethical review board of the Department of Psychology at the University of Sheffield (protocol number 053717).

**Table 1.** Full list of participants and the respective body regions where stretch measurements were taken. A checkmark (✓) indicates data that has been included in the paper. Data that has not been included is denoted by different letters; (v): successful data collection and processing, but the targeted body region is not fully visible or the speckle pattern does not extend across the region sufficiently, and therefore the data was excluded to ensure consistency across participants; (q): data quality was not sufficient and the digital image correlation process failed, for example because of smudging of the ink pattern or the presence of visible body hair in the targeted body region; (c): camera calibration issues prevented 3D reconstruction, for example due to inadvertent movement of the cameras between calibration and data collection.

|  | 0A | 0B | 01 | 02 | 03 | 04 | 05 | 06 | 07 | 21 | 22 | 23 | 24 | 25 | 26 |
| --- | --- | --- | --- | --- | --- | --- | --- | --- | --- | --- | --- | --- | --- | --- | --- |
| Wrist |  | (v) | (q) | (v) | ✓ | (v) | ✓ | (q) | ✓ |  |  |  |  |  |  |
| Knee | (q) |  | ✓ | (v) | (q) | (v) | ✓ |  |  |  |  | ✓ |  |  |  |
| Neck | (v) |  | ✓ | (v) | (q) | (v) |  |  |  | ✓ | ✓ | ✓ | ✓ | ✓ | ✓ |
| Back | (c) |  | (c) | (q) | (c) | (c) |  |  | ✓ |  |  |  |  |  | ✓ |
| Waist | ✓ |  | ✓ | (q) | ✓ | (q) |  |  |  |  |  |  |  |  |  |
| Face |  | ✓ | ✓ | (q) | ✓ | ✓ |  |  |  |  |  |  |  |  |  |

### Experimental protocol

At the start of the experimental session, ink-based random speckle patterns were applied to all included skin regions. Speckle patterns were generated in Speckle Generator 1.0.5 Build 209 (Correlated Solutions Inc., Irmo, USA) using a dot diameter of 1.27 mm, a dot density of 65%, and a dot variation of either 75% or 95%. Custom engraved rubber stamps (STAMPIT UK Ltd, London, UK) with these speckle patterns were manufactured in sizes 3.75 by 7.5 cm, 7.5 by 7.5 cm, and 14.75 by 14.75 cm, coated in water-based, dermatologically tested ink (Trodat 7011 black, Trodat GmbH, Wels, Austria) and lightly pressed on the participant’s skin. Depending on the size of the region tested, multiple stamps would be used. The participant then rested for 15-20 minutes to let the ink dry; in some instances warm air was directed to the skin through a hairdryer to help the drying process. In a subset of participants, speckle patterns were instead printed on A4 tattoo stencil paper using a thermal printer (LifeBasis, Shenzhen, China). Patches in the required sizes were then cut from these sheets and applied to the relevant skin regions using a small amount of stencil application solution (KMR distributions, Victorville, USA). In these cases, experiments could proceed without the rest period.

After the speckle patterns were applied, participants would sit, stand, or lie in front of the camera system, which was focused on the relevant skin region. A series of three (multi-camera) images was taken in the relaxed, unstretched pose, which served as the baseline for all stretch calculations. Participants would then assume one or more poses that were presumed to induce skin stretch. For the neck this was bending the head sideways while seated, for the waist bending the body sideways while standing, for the back curling into a fetal position, for the wrist flexing so the palm faced backwards, and for the knee flexing to different extents. For the face four different poses were included: opening the mouth widely (AU27, mouth stretch according to the Facial Action Coding System; Ekman and Friesen, 1976), puffing the cheeks (AU13, cheek puffer), pouting (AU22, lip funneler), and smiling (AU12, lip corner puller). Series of three images each were obtained in each pose. For both stretched and unstretched poses, the best (multi-camera) image from each series was selected for further processing.

### Image acquisition and processing

Images were acquired using a custom built setup that included four Arducam 64MP autofocus cameras (Arducam Technology Co. Ltd, Hong Kong) for synchronous operation connected via a dedicated UC-512 board (Uctronics, Nanjing, China) to a Raspberry Pi Model 4B (Raspberry Pi Ltd, Cambridge, UK). The board allowed taking 16MP images (4576×3472 pixels each) through all four cameras simultaneously. The cameras were mounted on a tripod using stereoscopic brackets, such that they formed the corners of a rough square of 20 by 20 cm. The setup was positioned at roughly 30 cm distance from the targeted skin region and supported by LED lights to provide illumination. The cameras were calibrated in pairs using standard stereo checkerboard images, resulting in reprojection errors of 3.3 pixels on average (across 17 individual camera calibrations), with a single pixel located centrally in the image covering a region of roughly 0.1 by 0.1 mm in size.

### Calculation of skin stretch measures

Images were processed using the open DuoDIC (Solav and Silverstein, 2022) package in Matlab (Mathworks Inc., Natick, USA). In short, using digital image correlation (DIC), the software matches small speckle patches both across cameras and across different poses by cross-correlating the images obtained from different cameras and at different poses under different image transformations. We typically used a patch radius of 60 pixels and a spacing of 10 pixels, yielding several thousand tracked points across the full skin patch at submillimeter resolution. In some instances, this process failed as reliable image correlations could not be found by the algorithm. This could happen because of deficiencies in the ink pattern (e.g. smudged ink), which prevented matching image patches. In other participants, body hair, for example on the arms or the face, impaired the correlation method, and shaving the targeted skin region might be required for robust tracking in such cases. 19 individual skin regions were ultimately successfully processed using DIC and further analysed (see Table 1 for the full list). First, by considering corresponding points in the images across multiple cameras, they were projected into 3D space and triangulated, yielding a three-dimensional surface. Tracked skin patches contained on average 9,327 triangular faces (range: 2,142-39,139), with an average size of 0.9 mm^2^ (0.5-2.4 mm^2^) each, covering an average skin area of 102.4 cm^2^, which could vary considerably depending on the region (21.2-397.5 cm^2^ from the face to the back, respectively). Next, by considering the difference in 3D shape between the relaxed pose and the target pose, the resulting stretch and compression along the two-dimensional surface was calculated for each triangle that made up the surface. Specifically, we used principal Lagrangian strains, which decomposes the deformation into values: the stretch along the direction where it is maximal (principal stretch) and the stretch or compression along the direction orthogonal to this principal direction.

For visualisations, we calculated the best-fitting plane for each skin patch and projected the surface on it, using custom code and the matGEOM Matlab package (Legland, 2023). Next, we resampled stretch magnitudes and orientations on a grid with 1 mm resolution. As the imaged regions were mostly flat, the vast majority of total summed strain for each body sites was constrained to this plane (97% on average, with a minimum of 90%).

### Validation of method and accuracy

To test the accuracy with which our setup could determine stretch under different viewing conditions, we systematically deformed the speckle patterns in Adobe Illustrator (Adobe Inc., San Jose, USA) and imaged print-outs of the patterns under different viewing angles. We then used the analysis pipeline described above to calculate the resulting deformations and stretch patterns, and compared them against the ground truth. We found excellent three-dimensional reconstruction accuracy with errors for individual tracked points of 0.2 mm on average for different viewing angles and under stretched conditions. Imaging the same baseline speckle pattern multiple times, either in the same view or under a changed viewing angle, should yield stretch values close to zero. Indeed, we found average absolute errors of 0.3% when the speckle patterns position was unchanged and 0.4% when the viewing angle was adjusted, demonstrating that inherent noise was low. When uni-axial uniform stretch of 33% was applied to the speckle pattern, the resulting stretch values differed on average by 0.9% in the same pose and 0.3% under a different viewing angle along the principal direction, and by 0.4% and 0.3%, respectively, in the orthogonal direction. Thus, errors were on average 1-2 orders of magnitude lower than the main effects reported in this study.

## ACKNOWLEDGEMENTS

MR was supported, in part, by a Sheffield Undergraduate Research Experience (SURE) fellowship. LDC was supported by a studentship from the MRC Discovery Medicine North (DiMeN) Doctoral Training Partnership (MR/N013840/1). HPS was supported by the Leverhulme Trust under Research Project Grant RPG-2022-031. We would like to thank Rory Holmes for help in developing the image acquisition software, Anika Kao, Greg Gerling, Sarah Crossland, and Claire Brockett for helpful suggestions improving the experimental setup, Chaona Chen for sharing knowledge about facial expressions, and all participants for supporting these experiments.

## AUTHOR CONTRIBUTIONS

MR, LDC, and HPS designed the research. MR, LDC, and HPS collected the data. MR and HPS performed the data analysis and prepared the figures. HPS wrote the manuscript, and MR and LDC contributed to review and editing.

## DATA AVAILABILITY

All processed DIC data and anonymized reference images are available on Zenodo (doi:10.5281/zenodo.13939433).

## Notes

### Competing Interest Statement

The authors have declared no competing interest.

